# Circulating biomarkers reflecting type III, IV and VI collagen remodeling are present in lung tissue of patients with pulmonary fibrosis and non-fibrotic controls

**DOI:** 10.64898/2026.02.10.704745

**Authors:** Helene Wallem Breisnes, Sissel Kronborg-White, Mathias Høj, Filipa Bica Simões, Diana Julie Leeming, Morten Asser Karsdal, Simon Francis Thomsen, Line Bille Madsen, Søren Helbo, Elisabeth Bendstrup, Jannie Marie Bülow Sand

## Abstract

**Background:** The extracellular matrix (ECM) is a dynamic network that provides structural support and maintains tissue homeostasis. Collagens are the main structural components of the ECM, occupying distinct tissue compartments and serving specialized roles. Dysregulated ECM remodeling involves an imbalance between collagen production and degradation, generating neoepitope-specific fragments that can be released into circulation. Serological measurements of these fragments can be used as biomarkers of disease and have been associated with progression and mortality in different fibrotic diseases, including pulmonary fibrosis (PF). This study aimed to investigate whether these systemic biomarkers originate from human lung tissue in patients with PF and non-fibrotic controls.

**Methods:** Lung tissue was collected from patients with PF (n = 21) and non-fibrotic controls (n = 21) and processed in parallel as formalin-fixed paraffin-embedded or snap-frozen samples. Serum samples were collected from patients with PF and healthy controls (n = 21). Neoepitope-specific biomarkers reflecting type III, IV, and VI collagen production (PRO-C3, PRO-C4, and PRO-C6) and degradation (C3M, C4M, C4Ma3, and C6M) were quantified in serum and proteolytically degraded lung tissue, and their spatial distribution was assessed by immunohistochemistry in lung tissue sections.

**Results:** All collagen remodeling biomarkers were significantly increased in serum of patients with PF compared with healthy controls (PRO-C3: p = 0.0006, all others: p < 0.0001). Collagen degradation fragments (C3M, C4M, and C6M) could be generated and released from both non-fibrotic and fibrotic human lung tissue following proteolytic cleavage with pepsin, collagenase, and/or MMP-9. All biomarkers were detected in lung tissue by immunohistochemical staining, with widespread distribution of type III and IV collagen fragments, whereas type VI collagen (PRO-C6) production showed a more compartment-specific pattern.

**Conclusions:** These findings demonstrated that neoepitope-specific collagen remodeling biomarkers, usually detected in circulation, are present and can be released from human lung tissue. Their spatial distribution suggests that ECM remodeling is heterogeneous and differs according to collagen type and distinct tissue compartments. Collectively, our findings support the use of collagen remodeling biomarkers as tools to assess ECM remodeling in pulmonary disease.

## Introduction

Interstitial lung disease (ILD) is an umbrella term for a wide range of inflammatory and fibrotic pulmonary diseases that impair gas exchange and lung function [1]. Idiopathic pulmonary fibrosis (IPF), one of the most severe and common ILDs, is characterized by progressive scarring of the lung interstitium, leading to worsening of dyspnea and, in some cases, a persistent dry cough, ultimately resulting in a reduced quality of life and respiratory failure [2]. IPF largely affects elderly adult males and has a median survival time of 3-5 years if left untreated [3]. Current clinical diagnosis relies largely on spirometry, imaging, and occasionally invasive tissue biopsies, highlighting the need for non-invasive biomarkers that can assist in early diagnosis, assess disease activity, progression, and response to therapy.

The extracellular matrix (ECM) is a dynamic and highly organized network of proteins that provides structural support, regulates cell behavior, and maintains tissue homeostasis. In the lung, ECM remodeling is a continuous and tightly regulated process; however, severe or repetitive injury can disrupt this balance, leading to dysregulated remodeling characterized by fibrosis, tissue stiffening, and impaired gas exchange [5]. Collagens are the major structural proteins of the ECM, and different collagen types occupy distinct ECM compartments, serving specialized roles. Within basement membranes, type IV collagen (COL4) forms a network that provides structural support to epithelial and endothelial cells and maintains tissue barrier integrity [6]. In the interstitial matrix, type III collagen (COL3) is a major structural component, forming dense fibrils that provide tensile strength and maintain tissue architecture [6]. In fibrotic conditions, COL3 is also the primary protein produced during early tissue remodeling [7]. At the interface between the basement membrane and interstitial matrix, type VI collagen (COL6) acts as a microfibrillar bridge, organizing the tissue architecture and mediating cell-matrix signaling [6].

Dysregulated ECM remodeling is characterized by an imbalance between collagen production and degradation. During remodeling, neoepitope-specific collagen fragments are released into circulation, where they can serve as biomarkers of tissue turnover [8]. These biomarkers have been widely studied across chronic diseases and have been associated with disease severity, progression, and mortality, including in IPF [9–15]. While elevated systemic levels of fragments reflecting COL3, COL4, and COL6 remodeling have been reported in IPF, it remains unclear whether these fragments originate from lung tissue. Here, we investigated whether circulating collagen remodeling biomarkers can be released from human lung tissue by combining tissue cleavage experiments and immunohistochemical analysis of non-fibrotic and fibrotic lungs, and by relating these tissue findings to biomarkers levels measured in serum.

## Materials and methods

### Study cohort

Patients who were scheduled to undergo transbronchial cryobiopsies as part of the diagnostic workup for fibrotic ILD at Aarhus University Hospital (AUH; n = 21) were invited to participate in the study. Lung tissue samples were obtained via bronchoscopy and subsequently formalin-fixed and paraffin-embedded, following standard protocols at AUH. Additionally, one tissue biopsy was frozen within 20-30 min after extraction and stored at -80 °C. Non-fibrotic control lung tissue was obtained from lung cancer patients (n = 21) undergoing surgical lung resection. The samples were taken from areas as distant as possible from the tumor-affected tissue. The non-fibrotic tissue was divided into separate samples that were either formalin-fixed and paraffin-embedded or frozen within 20-30 min after extraction and stored at -80 °C. For patients under investigation for ILD, blood samples were collected as part of the routine blood tests before the bronchoscopy. Briefly, blood samples were collected and gently mixed five times, before leaving to clot for 30-60 min at room temperature. The samples were then centrifuged at 2,000 g for 15 min at 4 °C and serum was stored within 30 min at -80 °C. As blood was not collected from non-fibrotic controls, serum from healthy controls (n = 21) was obtained from BioIVT (West Sussex, UK).

### Biomarker measurements

Neoepitope-specific biomarkers reflecting type III, IV, and VI collagen production (nordicPRO-C3^™^ [16], nordicPRO-C4^™^ [17], and nordicPRO-C6^™^ [18]) and MMP-mediated degradation (nordicC3M^™^ [19], nordicC4Ma3^™^ [20], and nordicC6M^™^ [21]) (all Nordic Bioscience, Herlev, Denmark) were quantified in serum from patients with PF (n = 18) from the AUH ILD cohort, as serum samples were not obtained from three patients, and healthy controls (n = 21). Samples were measured in duplicate, and values outside the measurement range were assigned the lower (LLMR) or upper limit of the measurement range (ULMR), as appropriate.

### Lung tissue cleavages

Frozen lung tissue samples from patients with PF or non-fibrotic controls were thawed, weighed, and homogenized in 0.01 M phosphate buffer saline (PBS, pH 7.4) at a ratio of 1 mg tissue to 3 μL PBS using a TissueLyser II (Qiagen, Hilden, Germany) at 30 Hz for 3 x 2 min with two 5 mm metal beads. Tissue lysates were then cleaved with pepsin or collagenase to allow for tissue exposure. Pepsin (cat. no. P7000, Sigma-Aldrich, St. Louis, MO, USA) was activated in 10 mM hydrochloric acid. Collagenase (cat. no. C9891, Sigma-Aldrich, St. Louis, MO, USA) was diluted 1:25 in matrix metalloproteinase (MMP) buffer (50 mM Tris-HCl, 200 mM NaCl, 10 mM CaCl_2_, 100 µM ZnCl_2_, 0.05% Brij-35, pH 7.5). Tissue lysates were pre-incubated with pepsin or collagenase (3:1 ratio) at 37 °C for 30 min. Following this, enzymes were inactivated by neutralizing pepsin with sodium hydroxide to pH ∼8 or heating collagenase-treated samples to 56 °C for 15 min. Following pre-treatment, recombinant pro-MMP-9 (cat. no. PF038, Merck, Darmstadt, Germany) was activated with 4-aminophenylmercuric acetate (14 mg/mL in dimethyl sulfoxide, 1:10 ratio) at 37 °C for 16-24 h. Samples were incubated for 3 h at 37 °C in MMP buffer, with or without addition of activated MMP-9 (final concentration 18.5 nM). Reactions were stopped by addition of ethylenediaminetetraacetic acid (40 μM), and samples were centrifuged at 13,000 g for 10 min at 4 °C. Supernatants were stored at -20 °C until analysis of collagen degradation biomarkers (nordicC3M^™^, nordicC4M^™^ [22], and nordicC6M^™^, all Nordic Bioscience). Cleavage efficiency was confirmed in parallel by SDS-PAGE analysis of carboxymethylated transferrin fragmentation (data not shown).

### Immunohistochemical staining

Formalin-fixed and paraffin-embedded lung tissue sections (6 μm) from patients with PF (n = 21) and non-fibrotic controls (n = 21) were used for IHC analysis. Sections were deparaffinized in toluene and rehydrated through graded ethanol (99%, 96%, 70%). Antigen retrieval was performed in 10 mM citrate buffer (pH 6) at 100 °C for 15 min, followed by washing in 0.01 M PBS. Endogenous peroxidase activity was blocked with 0.3% hydrogen peroxide for 30 min.

Primary antibodies against collagen neoepitopes reflecting production (nordicPRO-C3^™^, nordicPRO-C4^™^, and nordicPRO-C6^™^) and degradation (nordicC3M^™^ and nordicC6M^™^) were used. All neoepitopes were detected using monoclonal mouse anti-human antibodies (Nordic Bioscience) and diluted in 2% bovine serum albumin (BSA)-PBS: PRO-C3 (1:1000), PRO-C4 (1:400), PRO-C6 (1:2000), C3M (1:100), and C6M (1:100). Tissue sections were incubated with primary antibodies for 1 h at room temperature (PRO-C6, C6M) or 24 h at 4 °C (PRO-C3, PRO-C4, C3M).

For PRO-C3, PRO-C4, and C3M, an additional blocking step was performed for 30 min prior to antibody incubation (PRO-C3 and C3M: 2% rabbit serum; PRO-C4: 1% rabbit serum; all in 0.01 M PBS + 1% Tween-20). After washing, slides were incubated with a horseradish peroxidase-conjugated secondary antibody (polyclonal rabbit, anti-mouse, cat. no. 315-035-045, 1:100; Jackson ImmunoResearch, West Grove, PA, USA) prepared in 2% human serum + 2% BSA-PBS. Positive staining was developed using NovaRED for 10-20 min (cat. no. SK-4800, Vector Laboratories, Burlingame, CA, USA) and counterstained with hematoxylin for 2 min. Sections were dehydrated, mounted, and imaged with a 4x or 10x objective using Olympus BX60. Negative controls (no primary antibody) and absorption controls (antibody preincubated with synthetic peptide with the neoepitope antigen sequence) at 1:3 or 1:6 for 1 h at room temperature were included.

### Statistical analysis

All statistical analyses were performed using GraphPad Prism (version 10.5.0, San Diego, CA, USA). When comparing two groups, the Mann-Whitney test was used, and when comparing three or more groups, Kruskal-Wallis One-Way ANOVA with Dunn’s multiple comparisons test was applied. Spearman’s rank correlation test and chi-square test were used to investigate correlations and distributions. Patient characteristics were summarized as mean ± standard deviation (SD) or as count (percentage). Data are shown as scatter plots with median and interquartile range (IQR) indicated. Differences were considered statistically significant if p < 0.05. Asterisks indicate the following: * = p < 0.05, ** = p < 0.01, *** = p < 0.001, **** = p <0.0001, and ns = not significant.

## Results

### Collagen biomarkers are elevated in serum of patients with pulmonary fibrosis

Patients with PF were significantly older than healthy controls (70.6 [SD 7.3] vs. 47.4 [SD 13.1] years, p < 0.0001), while sex distribution was similar between groups (55.5% vs. 52.4% male, p = 0.843, **Table 2**).

**Table 1.** Patient characteristics of healthy controls and pulmonary fibrosis patients.

|  | Healthy controls (n = 21) | PF (n = 18) | p value |
| --- | --- | --- | --- |
| Age (years) | 47.4 ± 13.1 | 70.6 ± 7.3 | **< 0.0001** |
| Sex |  |  | 0.843 |
| Male | 11 (52.4%) | 10 (55.5%) |  |
| Female | 10 (47.6%) | 8 (44.4%) |  |
| BMI (kg/m<sup>2</sup>) | - | 29.1 ± 4.7 |  |
| FVC (% predicted) | - | 94.8 ± 15.8 |  |
| DLCO (% predicted) | - | 61.4 ± 13.6 |  |
| 6MWD (meters) | - | 474.6 ± 80.3 |  |

Data are shown as mean ± SD or n (%). Significant p values are highlighted in bold. BMI: body mass index, DLCO: diffusing capacity of the lungs for carbon monoxide, FVC: forced vital capacity, PF: pulmonary fibrosis, 6MWD: 6-minute walking distance.

**Table 2.** Baseline characteristics of non-fibrotic and pulmonary fibrosis patients.

|  | **Non-fibrotic (n = 21)** | **PF (n = 21)** | **p value** |
| --- | --- | --- | --- |
| Age (years) | 68.6 ± 8.1 | 69.6 ± 7.4 | 0.579 |
| Sex |  |  | 0.757 |
| Male | 11 (52.4%) | 12 (57.1%) |  |
| Female | 10 (47.6%) | 9 (42.9%) |  |
| BMI (kg/m<sup>2</sup>) | 25.5 ± 4.4 | 29.7 ± 4.9 | **0.010** |
| Smoking status |  |  | **0.046** |
| Smoker | 9 (42.6%) | 2 (9.5%) |  |
| Ex-smoker | 9 (42.6%) | 13 (61.9%) |  |
| Non-smoker | 3 (14.3%) | 6 (28.6%) |  |
| Packages pr. Year |  |  |  |
| Smoker | 49.2 ± 16.6 | 59 ± 22.6 | **0.014** |
| Ex-smoker | 38.6 ± 20.6 | 19 ± 12.1 | 0.764 |
| FVC (% predicted) | 114.9 ± 28.4 | 92.5 ± 16.9 | **0.006** |
| DLCO (% predicted) | 81.0 ± 17.3 | 59.1 ± 14.2 | **< 0.0001** |
| 6MWD (meters) | - | 484.1 ± 85.0 | - |

Data are shown as mean ± SD or n (%). Significant p values are highlighted in bold. BMI: body mass index, DLCO: diffusing capacity of the lungs for carbon monoxide, FVC: forced vital capacity, PF: pulmonary fibrosis, 6MWD: 6-minute walking distance.

All collagen remodeling biomarkers were significantly higher in serum from patients with PF compared with healthy controls (PRO-C3: p = 0.0006, all others: p < 0.0001, **Figure 1**). For both subject groups, none of the biomarkers correlated with age and for PF patients neither with BMI nor 6MWD (data not shown). Pulmonary function measurements correlated negatively with C4Ma3 (forced vital capacity [FVC]: p = 0.004, r = -0.647, diffusing capacity of the lungs for carbon monoxide [DLCO]: p = 0.002, r = -0.687) and C6M (DLCO: p = 0.007, r = -0.614) in patients with PF (**Figure 2**).

**Figure 1.**
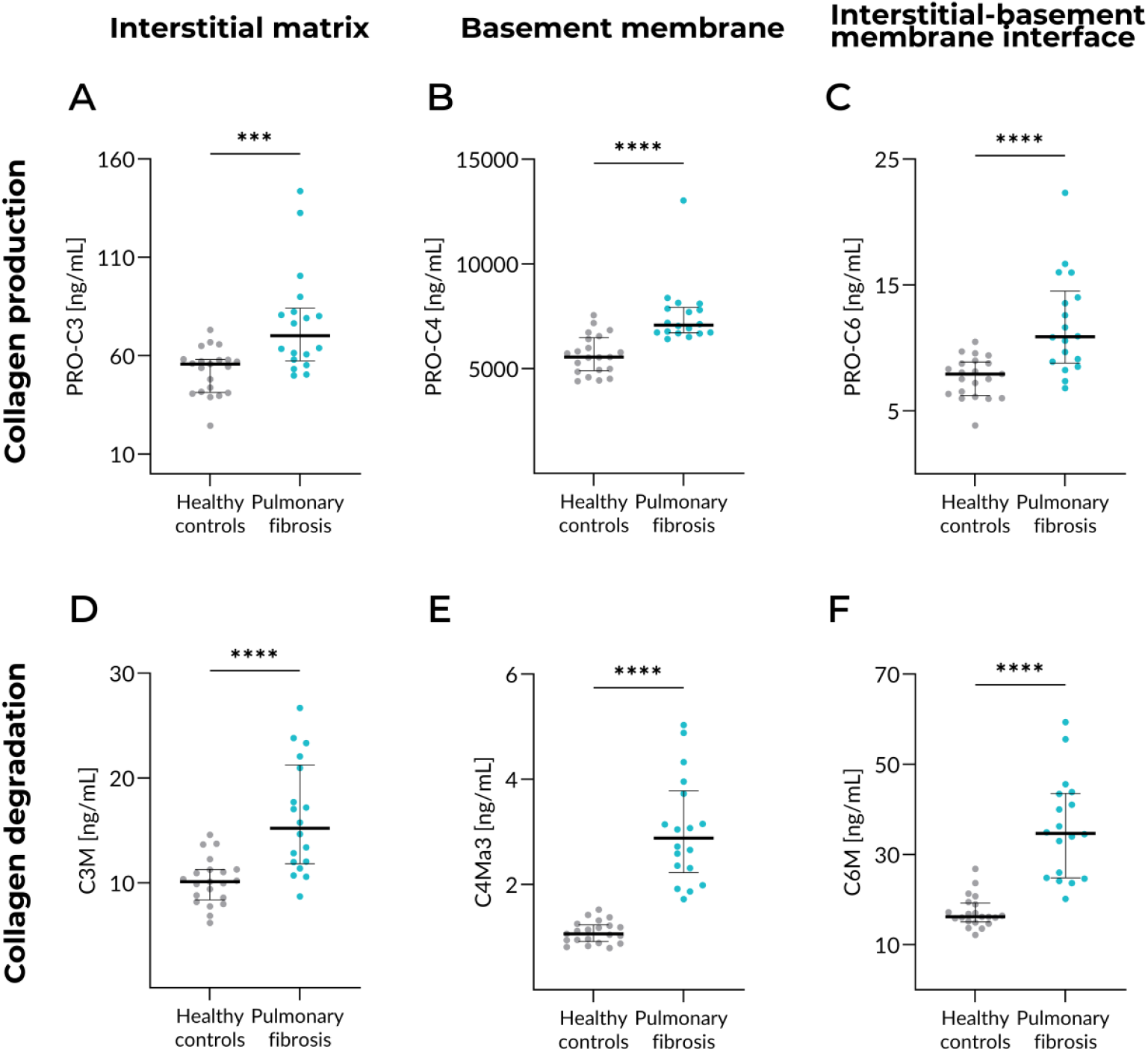
Collagen remodeling biomarkers were increased in serum of patients with pulmonary fibrosis. Biomarkers reflecting production of **A)** type III collagen (PRO-C3), **B)** type IV collagen (PRO-C4), and **C)** type VI collagen (PRO-C6) and degradation of **D)** type III collagen (C3M), **E)** type IV collagen (C4Ma3), and **F)** type VI collagen (C6M) were measured in healthy controls (n=21) and patients with pulmonary fibrosis (n=18). Type III collagen localizes in the interstitial matrix, type IV collagen localizes in the basement membrane, and type VI collagen localizes in the interface between the interstitial matrix and the basement membrane. Data were analyzed by Mann-Whitney test and are shown as scatter plots with median and interquartile range. PRO-C3: p = 0.0006, all others: p < 0.0001.

**Figure 2.**
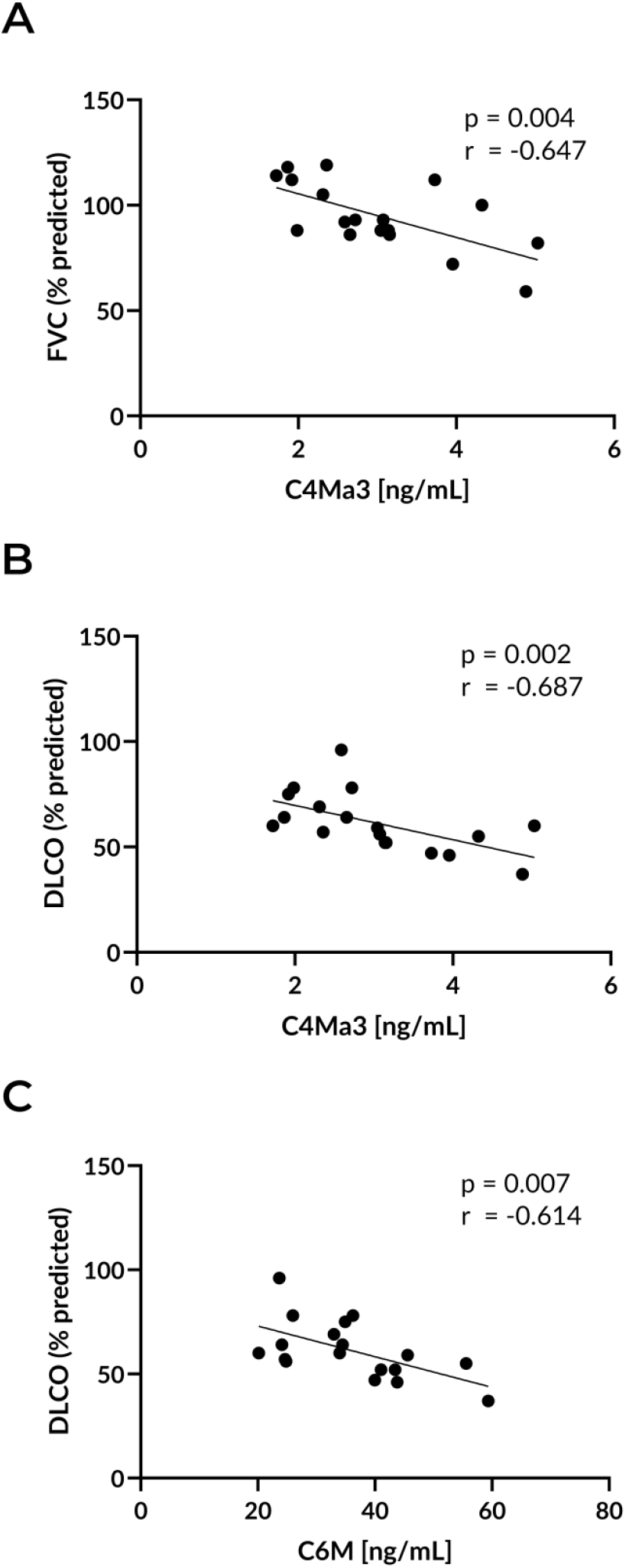
Correlation between serum C4Ma3 and C6M with FVC/DLCO in patients with pulmonary fibrosis. Correlation of the degradation fragment of type IV collagen (C4Ma3) with **A)** FVC (% predicted) and **B)** DLCO (% predicted). **C)** Correlation between degradation of type VI collagen (C6M) and DLCO (% predicted). n=18. Data were analyzed with Spearman correlation. DLCO: diffusing capacity of the lungs for carbon monoxide, FVC: forced vital capacity.

### Collagen degradation biomarkers originated from human lung tissue

The present study enrolled 42 participants in total, 21 non-fibrotic controls and 21 patients with PF, who were similar in age (68.6 [SD 8.1] vs. 69.6 [SD 7.4] years, p = 0.579) and sex distribution (52.4% vs. 57.1% male, p = 0.757). Patients with PF had higher BMI (29.7 [SD 4.9] vs. 25.5 [SD 4.4], p = 0.010) and a different smoking status, with fewer current smokers (9.5% vs. 42.6%) and more ex-smokers (61.9% vs. 42.6%, p = 0.046) than non-fibrotic controls. Pulmonary function was significantly reduced in patients with PF, with lower FVC (% predicted: 92.5 [SD 16.9] vs. 114.8 [SD 28.4], p = 0.006) and DLCO (% predicted: 59.1 [SD 14.2] vs. 81.0 [SD 17.3], p < 0.0001).

To investigate whether collagen degradation fragments originated from human lung tissue, we proteolytically cleaved the lung tissue using relevant proteases and measured biomarkers in the supernatants. The results showed that degradation fragments of type III, IV, and VI collagen could be generated and detected in human lung tissue following proteolytic cleavage (**Figure 3**). Pre-incubation with pepsin followed by MMP-9 cleavage resulted in a significant increase in C3M and C6M release compared with pepsin treatment alone, both in non-fibrotic and fibrotic tissue (all: p < 0.0001). Most concentrations exceeded the ULMR, and limited sample volume prevented dilution and remeasurement, making it unclear whether C3M and C6M levels differed between non-fibrotic controls and patients with PF. For COL4, C4M was generated by pre-incubation with collagenase and subsequent cleavage with MMP-9 when compared with collagenase alone (p < 0.0001) in non-fibrotic tissue, and a similar trend was seen in fibrotic tissue (p = 0.057). No difference was observed in C4M levels in non-fibrotic vs fibrotic tissue.

**Figure 3.**
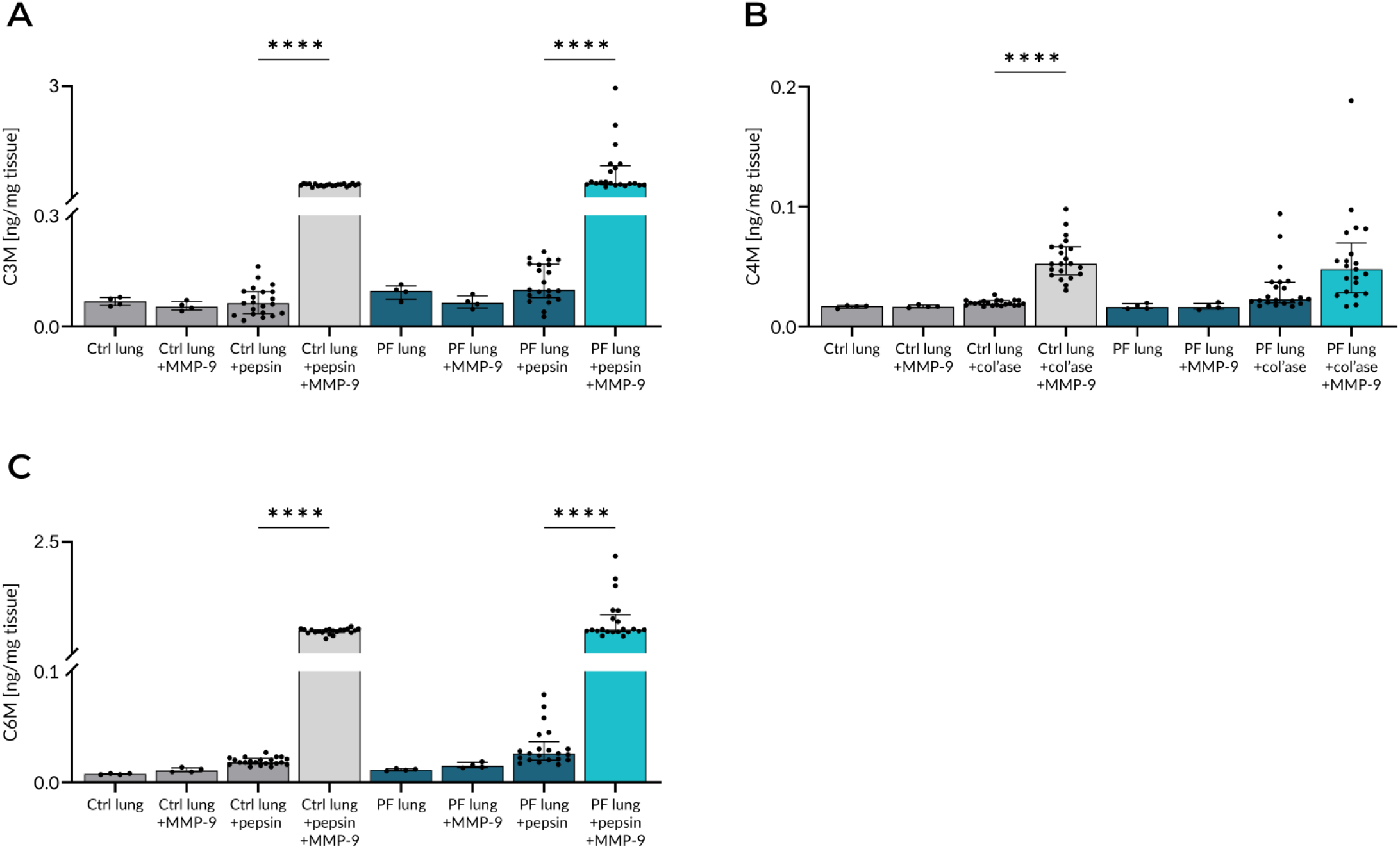
Collagen degradation fragments were detected in human lung tissue supernatants after proteolytic digestion. Human lung tissue from patients with PF and non-fibrotic controls was proteolytically digested with MMP-9, pepsin, collagenase, or a combination of those, and biomarkers reflecting **A)** type III collagen degradation (C3M), **B)** type IV collagen degradation (C4M), and **C)** type VI collagen degradation (C6M) were measured in supernatants. Biomarker concentrations were normalized to the tissue weight. Data were analyzed by Kruskal-Wallis test. Asterisks indicate: **** p < 0.0001. Col’ase: collagenase, ctrl: control, MMP: matrix metalloprotease, PF: pulmonary fibrosis.

### Collagen fragments were present in human lung tissue

Immunohistochemical staining of collagen remodeling fragments revealed that all investigated collagen fragment types were present in human lung tissue. Both COL3 production (PRO-C3) and degradation (C3M) fragments were widely distributed throughout the entire interstitium, around airways and blood vessels, and adjacent to epithelial layers (**Figure 4**), and both fragments appeared to stain more intensely in fibrotic compared to non-fibrotic tissue. Staining of the COL4 production fragment (PRO-C4) showed strong, widespread distribution throughout both non-fibrotic and fibrotic tissue (**Figure 5**), and the staining appeared stronger in fibrotic tissue. Staining of the COL6 production fragment (PRO-C6) localized around airways and blood vessels in both non-fibrotic and fibrotic tissue, as well as in the parenchyma of fibrotic interstitial regions in PF tissue (**Figure 6A**). The COL6 degradation fragment (C6M) was distributed throughout the entire tissue of both non-fibrotic and fibrotic lungs (**Figure 6B**). No differences were observed in PRO-C6 and C6M staining intensities in non-fibrotic versus fibrotic lung tissue.

**Figure 4.**
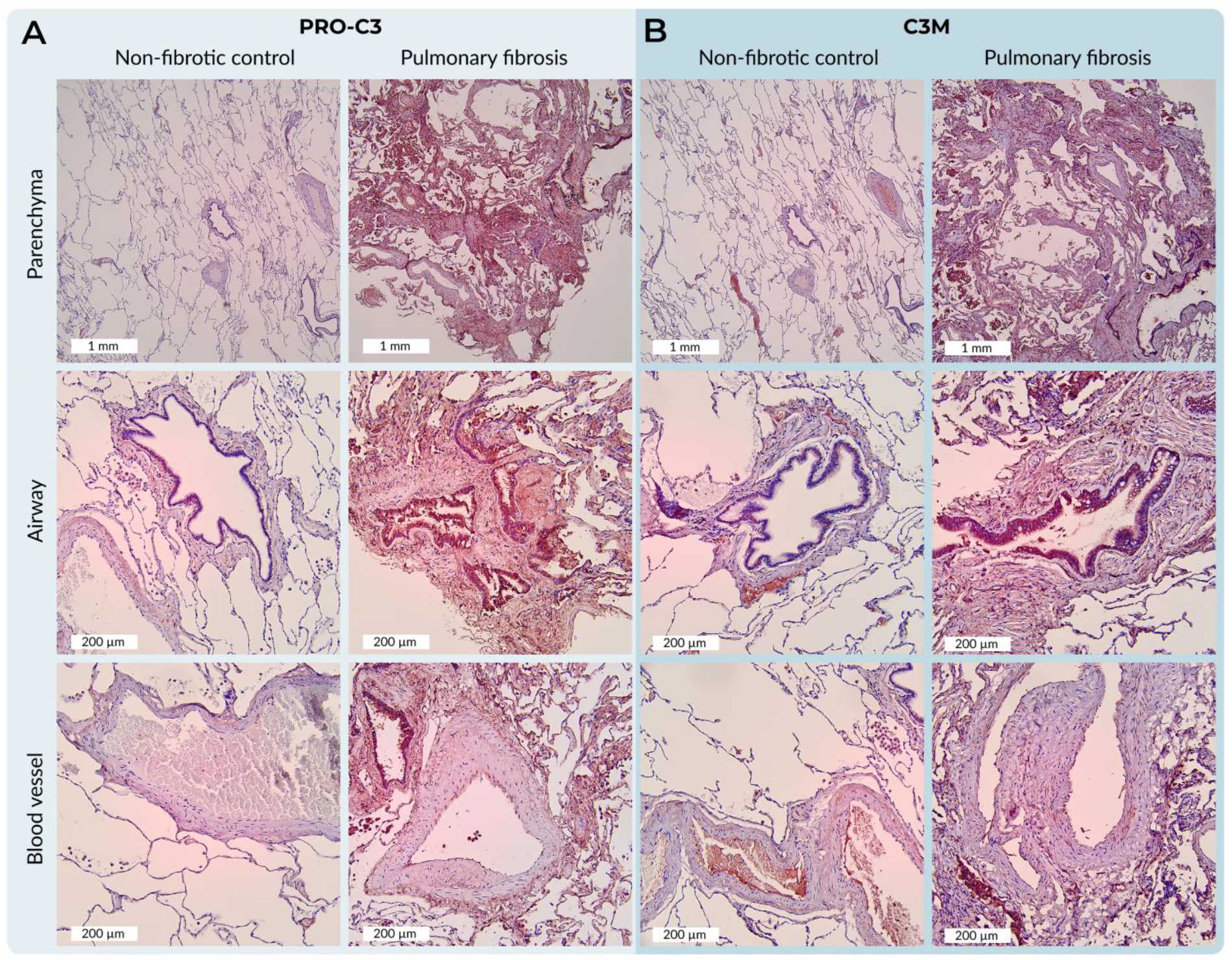
Type III collagen remodeling fragments could be detected in human lung tissue. Non-fibrotic controls (n=21) and PF patients (n=21) were stained using antibodies targeting fragments reflecting **A)** type III collagen production (PRO-C3) and **B)** type III collagen degradation (C3M). Images in the top row are of parenchyma and have a scale bar of 1 mm. The middle row shows representative images of airways and the bottom row of blood vessels, both with a scale bar of 200 µm. Red/brown staining represents NovaRed detection of the target antibody, while blue staining corresponds to hematoxylin counterstaining. PF: pulmonary fibrosis.

**Figure 5.**
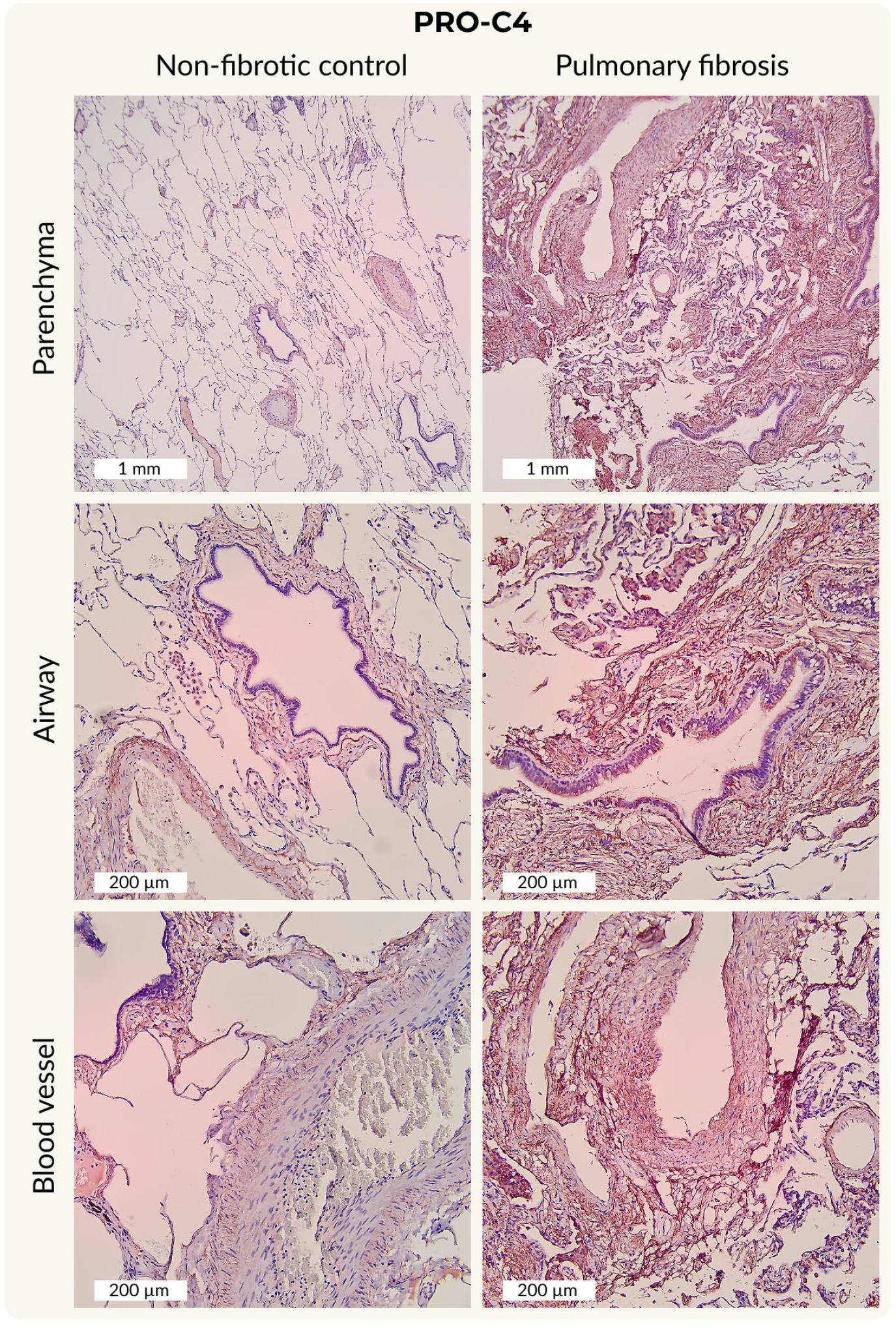
Type IV collagen formation fragments could be detected in human lung tissue. Tissue from non-fibrotic controls (n=21) and patients with PF (n=21) were stained with antibodies targeting a fragment of type IV collagen reflecting its production (PRO-C4). The images in the top row are of parenchyma and have a scale bar of 1 mm. The middle row shows representative images of airways and the bottom row of blood vessels, both with a scale bar of 200 µm. Red/brown staining represents NovaRed detection of the target antibody, while blue staining corresponds to hematoxylin counterstaining. PF: pulmonary fibrosis.

**Figure 6.**
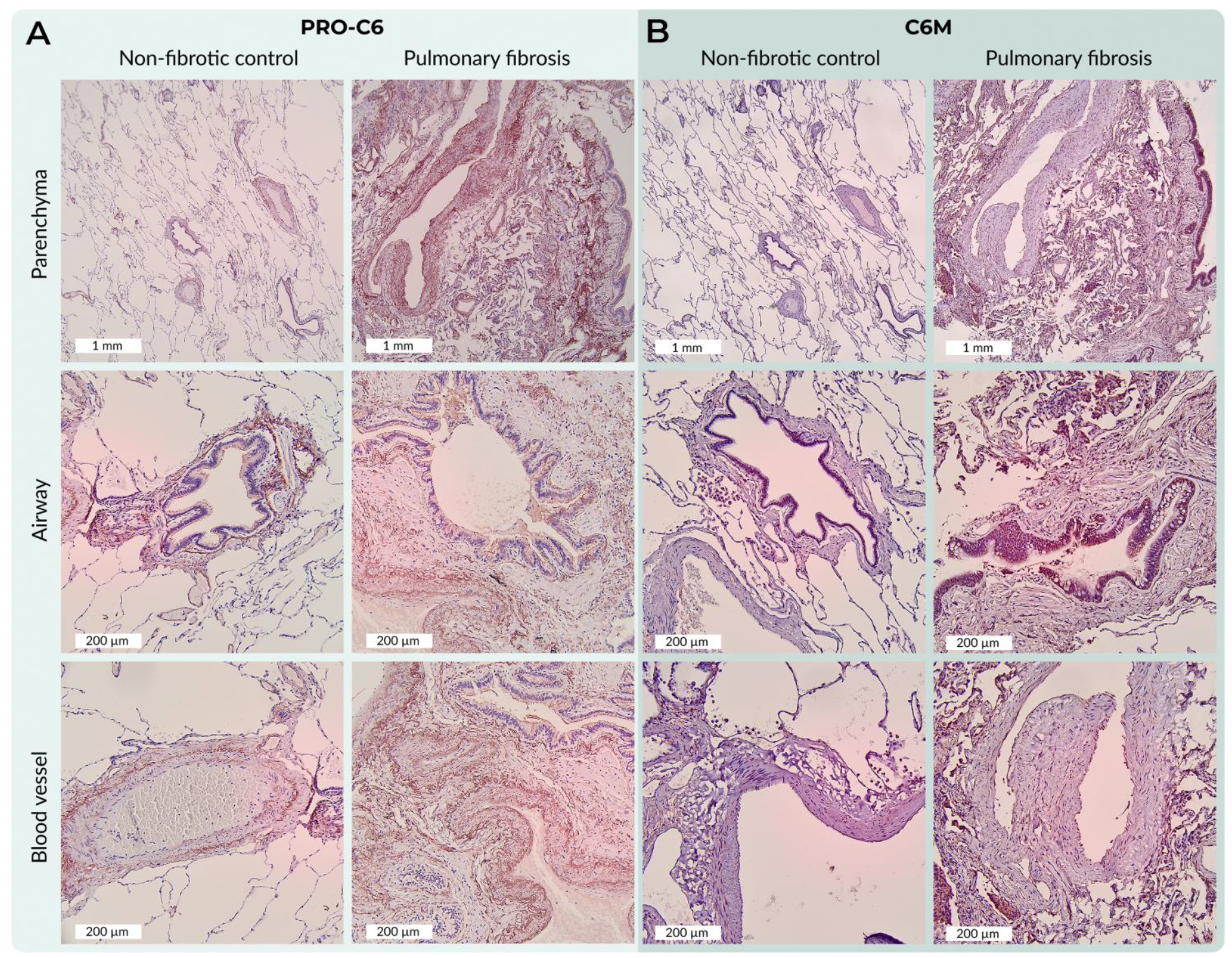
Type VI collagen remodeling fragments could be detected in human lung tissue. Tissue from non-fibrotic controls (n=21) and patients with PF (n=21) were stained with antibodies targeting fragments reflecting **A)** type VI collagen production (PRO-C6) and **B)** type VI collagen degradation (C6M). The images in the top row are of parenchyma and have a scale bar of 1 mm. The middle row shows representative images of airways and the bottom row of blood vessels, both with a scale bar of 200 µm. Red/brown staining represents NovaRed detection of the target antibody, while blue staining corresponds to hematoxylin counterstaining. PF: pulmonary fibrosis.

## Discussion

This study demonstrated that neoepitope-specific fragments of COL3, COL4, and COL6 remodeling can be detected in serum, are present and can be released from human lung tissue, and can be visualized *in situ*, providing a comprehensive tool to understand ECM remodeling in fibrotic and non-fibrotic lungs.

Neoepitope-specific collagen remodeling biomarkers have previously been shown to be elevated systemically in patients with IPF and other fibrotic diseases [9–13] and to associate with disease severity, progression, and mortality, and to be modulated by treatment [14, 15, 23, 24]. In this study, we sought to validate these observations in our cohort of patients with newly diagnosed PF. Consistent with previous reports, all evaluated biomarkers of collagen production and degradation were significantly increased in patients with PF compared with healthy controls. These elevations were observed across biomarkers reflecting the interstitial matrix, basement membrane, and the interstitial–basement membrane interface, highlighting that multiple structural compartments of the lung ECM are affected in PF.

Tissue cleavage experiments showed that proteolytic degradation of human lung tissue generated distinct collagen fragments that were detectable with neoepitope-specific biomarkers. Pepsin pre-treatment facilitated the generation of the specific type III (C3M) and VI (C6M) collagen degradation fragments, whereas the generation of a type IV (C4M) collagen degradation fragment required pre-treatment with collagenase. Collagenase facilitates complete degradation of the collagen triple helix, whereas pepsin mainly removes the non-collagenous sections while leaving the triple helix intact [25, 26]. As COL4 is structurally distinct from interstitial matrix collagens and embedded in a dense, cross-linked network, this localization may limit protease accessibility, potentially requiring more extensive proteolytic digestion for C4M to be released from the tissue. In the development of the C4M assay, only MMP-9 was used to cleave COL4 [22]. However, that experiment used purified COL4 from human placenta, whereas this study used native human lung tissue biopsies, highlighting the importance of protease accessibility. Collectively, these findings suggest that COL3, COL4, and COL6 degradation fragments are present and can be released from both non-fibrotic and fibrotic human lung tissue.

COL3 and COL6 have previously been shown to be increased in fibrotic tissue [27]. In this present study, the corresponding degradation fragments, C3M and C6M, were increased in the blood of patients with PF compared with healthy controls. Tissue measurements, however, could not be fully evaluated as most concentrations exceeded the ULMR, making it unclear whether disease-related differences in tissue concentrations of C3M and C6M exist between non-fibrotic and fibrotic tissue. For C4M, no significant difference in tissue concentrations was observed between patients with PF and non-fibrotic controls. Based on the accumulation and remodeling of basement membrane structures in fibrotic lung disease, one might expect more COL4 in fibrotic tissue. Indeed, COL4 staining in a recent study demonstrated increased COL4 deposition in IPF compared with non-fibrotic lung, followed by a reduction in regions with dense fibrosis, reflecting progressive basement membrane disruption [28]. The lack in difference in C4M levels in fibrotic and non-fibrotic tissue supernatants may reflect extensive proteolytic degradation, resulting in degradation of the C4M neoepitope. In addition, due to differences in tissue sample sizes, biomarker concentrations were normalized to tissue weight. This normalization could limit the interpretation of the absolute levels of degradation fragments, particularly given that fibrotic tissue has an increased matrix density and altered composition [29]. As a result, the normalized biomarker levels may not accurately reflect true differences in total fragment abundance between non-fibrotic and fibrotic tissue and may explain the difference to serum biomarker levels which were increased in patients with PF compared with healthy controls.

Immunohistochemical staining confirmed the presence of collagen remodeling fragments within human lung tissue. COL3 and COL4 fragments were broadly distributed throughout the tissue and appeared to stain more intensely in fibrotic compared with non-fibrotic tissue. The observed stronger staining likely reflects a higher local expression of COL3 and COL4 and their respective remodeling fragments within fibrotic tissue, suggesting active collagen remodeling in fibrotic regions. The COL6 degradation fragment (C6M) showed similar widespread distribution throughout the tissue, however, the COL6 production fragment (PRO-C6) exhibited more compartment-specific patterns, tending to localize mainly around airways and blood vessels in both non-fibrotic and fibrotic tissue, in addition to heterogeneous staining in the parenchyma of fibrotic interstitial regions. These observations are consistent with a recent study, which showed that PRO-C6 staining localized to the same regions in lungs of controls and severe IPF patients [30]. That study also quantified positively stained tissue and found that PRO-C6 expression was lower around airways, while C6M was lower in all tissue regions in IPF compared with non-fibrotic controls, suggesting that IPF lungs were characterized by region-specific alterations in COL6 remodeling. Considering the widespread distribution of COL3 and COL4 fragments, in contrast to the more compartment-specific patterns of COL6, it is likely that pathological ECM remodeling is unique for different collagens and matrix compartments. The increased systemic biomarker levels indicate that, while some ECM fragments are retained within the tissue, a substantial proportion is released into circulation, supporting their use as tools for non-invasive measurements.

While this study provides valuable insights into the clinical relevance of evaluating circulating collagen remodeling biomarkers in patients with PF, it is important to consider certain limitations. Circulating biomarker levels in non-fibrotic controls were assessed using healthy control serum rather than blood samples collected in parallel with tissue procurement. Although matched tissue and blood samples from non-fibrotic individuals would have been ideal, this was not feasible. In addition, healthy controls differed in age from patients with PF; however, correlation analyses revealed no significant associations between age and any of the evaluated biomarkers. Furthermore, technical constraints limited comprehensive quantification of all fragments released from degraded lung tissue, limiting the ability to fully assess disease-related differences in tissue-derived fragment abundance, and quantitative assessment of collagen fragment staining and discrimination between lung tissue compartments were not possible. These aspects warrant further investigation in future studies.

Taken together, this study demonstrates that neoepitope-specific collagen remodeling biomarkers, which are systemically elevated in patients with PF, are present and can be released from both non-fibrotic and fibrotic human lung tissue. Furthermore, their spatial distribution within human lung tissue suggests that ECM remodeling is heterogeneous and differs across collagen types and distinct tissue compartments. Collectively, our findings support the use of these biomarkers as tools to assess ECM remodeling in pulmonary disease.

## Abbreviations

6MWD: 6-minute walking distance
BMI: body mass index
BSA: bovine serum albumin
C3M: type III collagen degradation
C4M: type IV collagen degradation
C4Ma3: type IV collagen a3 chain degradation
C6M: type VI collagen degradation
DLCO: diffusing capacity of the lungs for carbon monoxide
ECM: extracellular matrix
FVC: forced vital capacity
ILD: interstitial lung disease
IPF: idiopathic pulmonary fibrosis
MMP: matrix metalloproteinase
PBS: phosphate buffered saline
PF: pulmonary fibrosis
PRO-C3: type III collagen production
PRO-C4: type IV collagen production
PRO-C6: type VI collagen production
SD: standard deviation

## Declarations

### Ethics approval and consent to participate

This study was approved by the Regional Ethics Committee (H-22051172) in Denmark and conducted in accordance with the Declaration of Helsinki. The healthy samples were collected in compliance with the REC recommendations according to the qualified vendor BioIVT,

### Consent for publication

Not applicable

### Availability of data and materials

The datasets used and/or analyzed during the current study are available from the corresponding author on reasonable request.

### Competing interests

FBS, DJL, MAK, and JMBS are employees and may be shareholders of Nordic Bioscience, a company involved in discovery and development of biochemical biomarkers. HWB is a PhD student and MH was a MSc student at the University of Copenhagen conducting their project in collaboration with Nordic Bioscience. The other authors have no conflicts of interest related to this study.

### Funding

This study was funded by the Danish Research Foundation.

### Authors’ contributions

Study concept and design: HWB, SKW, EB, JMBS. Acquisition of data: HWB, SKW, MH, LBM, SH, EB. Analysis and interpretation of data: HWB, MH, FBS, JMBS. Drafting of the manuscript: HWB. Critical revision of the manuscript for important intellectual content: SKW, FBS, DJL, MAK, SFT, EB, JMBS. All authors read and approved the final manuscript.

## Acknowledgements

We wish to acknowledge all the people involved in this project. We thank all the study participants and the medical, nursing, and technical staff who were responsible for collecting the samples for the study. We are also grateful to the organization that funded the study.

## Notes

### Competing Interest Statement

F.B. Simoes, M.A. Karsdal, D.J. Leeming, and J.M.B. Sand are employees and may be shareholders of Nordic Bioscience, a company involved in discovery and development of biochemical biomarkers. H. W. Breisnes is a PhD student and M. Hoej was a MSc student at the University of Copenhagen conducting their project in collaboration with Nordic Bioscience. The other authors have no conflicts of interest related to this study.

